# No detectable effect of P2 neurons on female post-mating changes in sleep and feeding in *Drosophila melanogaster*

**DOI:** 10.64898/2026.09.10.750755

**Authors:** Sushant Potdar, Tasnim Makhmudova, Ava Schenck-Davis, Sofie Y. N. Delbare, Asha M. Jain, Yassi Hafezi, Andrew G. Clark, Mariana F. Wolfner

## Abstract

In many insects, females exhibit behavioral changes after mating. This plasticity can be mediated by interactions between male-derived seminal fluid proteins (SFPs) and the female nervous system. The *Drosophila melanogaster* seminal fluid protein Sex Peptide (SP) causes mated females to sleep less during the day and consume more food than unmated females, among other behavioral changes. Although the SP-sensing neural circuits that convey information to the female’s brain are known, the downstream neurons that affect post-mating behaviors remain unknown. P2 neurons in the fly’s central brain are good candidates for regulating post-mating changes in sleep and food consumption, as they express feeding- and sleep-responsive neuropeptide F (NPF), directly innervate the fly’s sleep center (the dorsal fan-shaped body, dFB), and are responsible for social-isolation-induced changes in sleep and food consumption. Using the GAL4/UAS system, we activated or silenced P2 neurons in mated and unmated females. We measured sleep duration using Drosophila Activity Monitors (DAM) and food consumption using Capillary Feeder (CAFE) and blue-dye assays. We found that P2-neurons are not responsible for post-mating changes in sleep and food consumption: mated females slept less during the day and ate more compared to unmated females irrespective of P2 neuron perturbation. Together, these results indicate that P2 neurons are unlikely to be primary downstream neurons through which male-transferred SP regulates post-mating sleep and food consumption. Future studies could determine whether P2 neurons regulate other post-mating behaviors or modulate sleep and feeding behaviors in a context-dependent manner.

**Article summary:** Post-mating female *Drosophila melanogaster* sleep less and consume more food, initiated by male-transferred Sex Peptide. But the neurons involved in these post-mating behavioral changes are unknown. P2 neurons may modulate post-mating sleep and food consumption, similar to their known role in a social-isolation context. Ectopic activation and silencing of P2 neurons did not affect post-mating female sleep and food consumption: mated females slept less and consumed more food than unmated females. This study eliminates P2 neurons as targets of female post-mating sleep and feeding, thereby opening new avenues to investigate the roles of other neurons in modulating these post-mating behaviors.

## Introduction

Many animals change their behavior after mating as their priorities shift from survival to reproduction and, in some cases, to care for offspring (Wedell 2005; Wigby et al. 2020; Hopkins and Perry 2022; Guan et al. 2023). Males decrease aggression and courtship and may cease mating to replenish sperm and seminal fluid resources for future matings (Sun et al. 2026), whereas females invest more time and resources in egg production and laying (Wang et al. 2020), avoid remating (Dougherty 2024), and prioritize parental care (Rogers and Bales 2019). This behavioral plasticity is mediated, at least in part, through changes in the brain (reviewed in Afkhami 2024; Sun et al. 2026). Here, we explored the potential role of specific brain neurons in mediating mating-induced changes in female behaviors using the *Drosophila melanogaster* model.

*D. melanogaster* females undergo many behavioral changes after mating, collectively termed post-mating responses. Females lay more eggs (Chen et al. 1988; Aigaki et al. 1991; Afkhami 2024), reduce sexual receptivity (Manning 1967; Chapman et al. 2003; Wang et al. 2021), sleep less, especially during the day (Isaac et al. 2010; Garbe et al. 2016), eat more food (Carvalho et al. 2006), and change their dietary preferences (Walker et al. 2015; Liu et al. 2024). These post-mating behavioral changes are initiated by the male-transferred seminal fluid protein Sex Peptide (SP). SP binds to the sex peptide receptors (SPR) in the sensory neurons of the female reproductive tract.

These neurons signal the receipt of SP to the pC1 neurons in the female brain via neurons in the abdominal ganglion (Yapici et al. 2008; Häsemeyer et al. 2009; Yang et al. 2009; Wang et al. 2020; Wang et al. 2021; Laturney et al. 2023). pC1 neurons control downstream circuits that regulate multiple post-mating behavioral shifts. For example, pC1 neurons activate downstream oviposition neurons (oviINs, oviDNs, and oviENs; Wang et al. 2020; Afkhami 2024), vaginal plate opening (vpo) neurons (Wang et al. 2021), and Lgr3+ neurons (Laturney et al. 2023), which regulate female post-mating egg-laying, sexual receptivity, and sugar intake, respectively. However, the neurons that mediate post-mating changes in daytime sleep and overall food consumption are unknown.

One set of neurons in the fly central brain that could potentially regulate these female post-mating behaviors are the P2 neurons. P2 neurons express the *Drosophila* neuropeptide F (NPF) which regulates sleep and feeding (Chung et al. 2017; Shao et al. 2017; Li et al. 2021). Moreover, P2 neurons innervate the dorsal fan-shaped body (dFB) of the fly’s central brain, which regulates sleep-wake cycles in a nutrition-dependent manner (Duhart et al. 2023). On a sucrose-only food medium that reduces egg laying, thereby reducing larval interference in activity assays (Yang et al. 2008; Catterson et al. 2010; Yang et al. 2015; Garbe et al. 2016), mated females reduced their daytime but not nighttime sleep (Duhart et al. 2023). However, when sleep was measured on standard food containing yeast, mated females also reduced their nighttime sleep (Duhart et al. 2023). Silencing dFB neurons eliminated the effects of mating on nighttime sleep in females on standard, but not sucrose-only food (Duhart et al. 2023).

P2 neurons are also implicated in social isolation-induced reduction in daytime sleep and increased feeding (Li et al. 2021), phenotypes that resemble female post-mating responses (Carvalho et al. 2006; Isaac et al. 2010; Laturney et al. 2023). Silencing P2 neurons in chronically socially-isolated flies (7d) reversed the effects of social isolation: those flies slept and ate similarly to group-reared flies. Conversely, activation of P2 neurons in acutely isolated flies (1d) increased feeding and decreased daytime sleep (Li et al. 2021). Whether NPF-expressing P2 neurons that innervate the dFB regulate mated female’s daytime or nighttime sleep, or food intake remains unknown.

Intrigued by potential parallels to the social isolation findings, we asked if P2 neurons modulate reduced sleep and increased feeding in females after mating (Garbe et al. 2016; Li et al. 2021; Duhart et al. 2023). We hypothesize that if P2 neurons induce post-mating changes in sleep and feeding, then activation of P2 neurons in unmated females would reduce their daytime sleep and increase their food consumption to the levels observed in mated females. Conversely, silencing P2 neurons in mated females would increase their daytime sleep and reduce their food consumption to the levels observed in unmated females. However, we found that activation or silencing of P2 neurons did not impact post-mating changes in female sleep and feeding. Thus, other neurons either downstream of pC1 neurons, or independent of the pC1 neuronal pathway, must regulate these post-mating female behaviors.

## Results

### Manipulating P2 neurons did not modulate female post-mating decrease in daytime sleep

At 29°C on sucrose-only food medium, mated females of all genotypes (*P2>dTRPA1*, *P2>Kir2.1* and their respective controls) slept less than unmated females (ANOVA: *F*_1,261_ = 92.9, *p* < 2e-16; Figure 1a, b, e, f, I). This reduction occurred primarily during the day (ANOVA: *F*_1,261_ = 71.8, *p* = 1.7e-15; Figure 1j), with only brief reduction in sleep at night (ANOVA: *F*_1,261_ = 63.1, *p* = 5.8e-14; Figure 1k).

**Figure 1:**
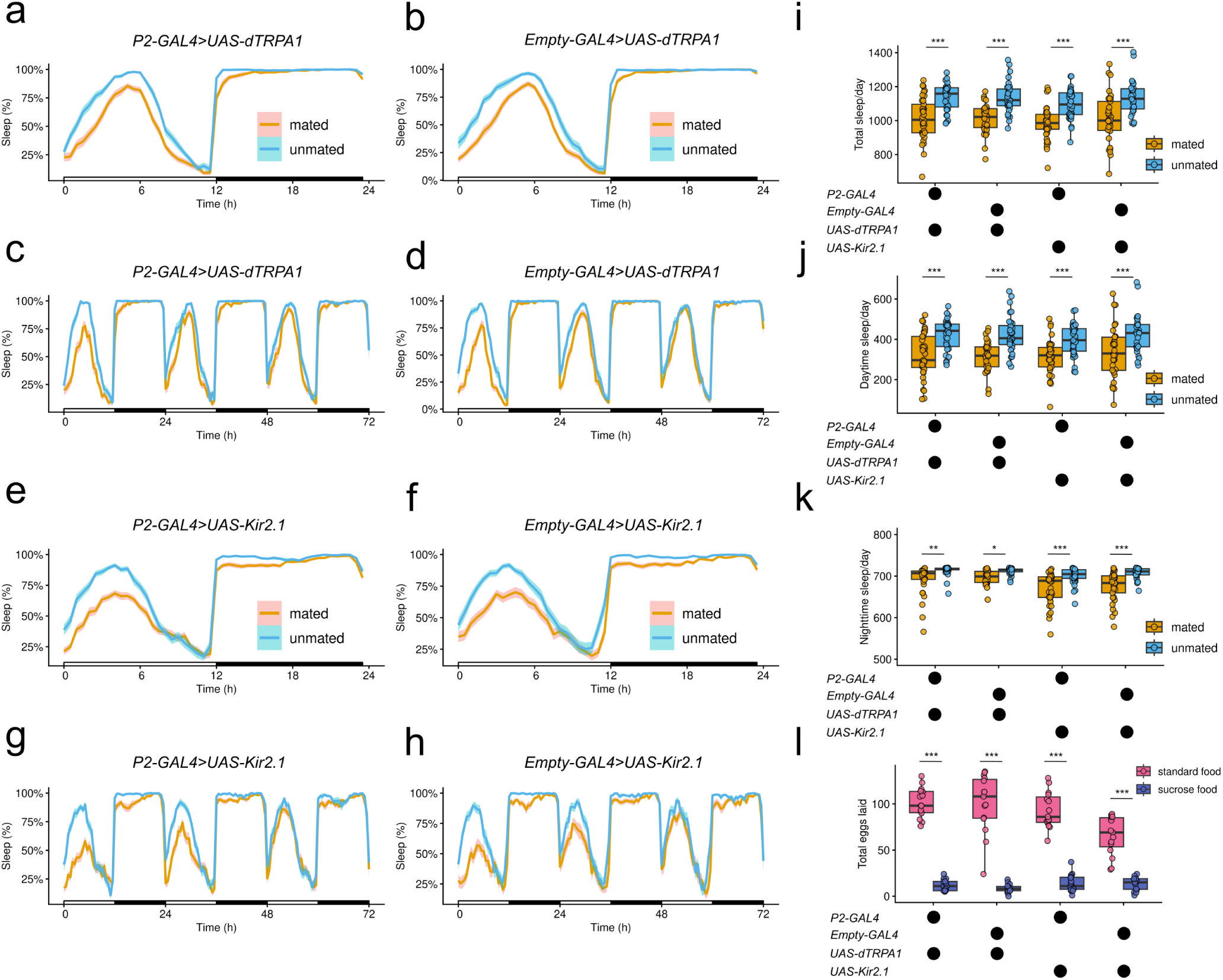
P2 neuron perturbation does not affect female post-mating sleep. Sleep profiles averaged across 3 L:D cycles in a) P2 neuron activated mated (*n*=37) and unmated (*n*=33) females, and its b) no GAL4 control (*EM>dTRPA1*) mated (*n*=33) and unmated (*n*=32) females, (e) P2 neuron silenced mated (*n*=37) and unmated (*n*=36) females, and its (f) no GAL4 control (*EM>Kir2.1*) mated (*n*=33) and unmated (*n*=31) females. (c), (d), (g), and (h) depict daily sleep rhythms of the same genotypes and mating status as in (a),(b), (e), and (f) respectively. Sleep for all flies was measured at 29°C on sucrose-only food. Mated females of all genotypes slept less compared to unmated females, averaged across L:D cycles, as measured by (I) total sleep (min); (j) daytime sleep (min); and (k) nighttime sleep (min). (l) Mated females laid more eggs on standard food (n=15/genotype) compared to sucrose-only food (n=15/genotype) averaged across three days of egg-laying. Sleep and egg-laying were statistically compared using two-way ANOVA (sleep = parametric, egg laying = aligned rank transform) followed by Sidak’s/Tukey’s posthoc pairwise comparison. All boxplots depict median, 25^th^ and 75^th^ percentiles, and whiskers represent 1.5X inter-quartile range. Dots in box and whisker plots represent sampled individuals. \**p*<0.05, \*\**p*<0.01, \*\*\**p*<0.001

However, unlike the ∼7 day effect of SP on female daytime sleep (Isaac et al. 2010), in our study mated females of all these genotypes reverted to an unmated-like sleep state by day 3 (Figure 1c, d, g, h). We hypothesized that the reversion to unmated sleep levels on sucrose-only food could be due to lack of energy expenditure, including due to egg-laying on that substrate, as mated females of all genotypes laid more eggs on the standard glucose-yeast food than sucrose-only food (ANOVA: *F*_1,112_ = 304.5, *p* < 2.22e-16; Figure 1l), every day across 3 consecutive days (ANOVA Food*Day*Genotype: *F*_1,336_ = 2.15, *p* < 0.05; Supplementary Figure 2). We also conducted the sleep assays at 25°C on *P2>Kir2.1, EM>Kir2.1*, and *CS* females. Here too, post-mating sleep reduction reverted before 7d (in this case, at 4d; Supplementary Figure 3b, d, f, g, h, l). All mated females with the *UAS-Kir2.1* construct (control and experimental) also slept less during the night compared to their unmated counterparts. This effect was not seen in mated *CS* females (ANOVA genotype*mating status: *F*_2,170_ = 6.5, *p* = 0.001; Supplementary Figure 3a, b), suggesting that genetic-background may influence the effect of mating on nighttime sleep.

### Manipulating P2 neurons did not modulate female post-mating increase in food consumption

We next used CAFÉ assays to compare food consumption for 24 hours by mated and unmated females with activated P2 neurons (*P2>dTRPA1;* 29°C), silenced P2 neurons (*P2>Kir2.1;* 25°C), and their respective controls. Mated females of all genotypes consumed more food than their unmated female counterparts (ANOVA: *F*_1,100_ = 63.17, *p* < 2.94 × 10^-12^; Figure 2a). Using the blue dye-based assay, we also found that mated females with silenced P2 neurons (*P2>Kir2.1*) and their matched controls consumed more food in a 1-hr period than their unmated counterparts (*P2>Kir2.1*: *t* = 7.23, df = 547, p < 0.0001; *EM>Kir2.1: t* = 5.64, df = 547, *p* < 0.0001; Figure 2b). This result was consistent across four trials (Supplementary Figure 4). Overall, our results suggest that manipulating P2 neurons did not influence female post-mating food consumption.

**Figure 2:**
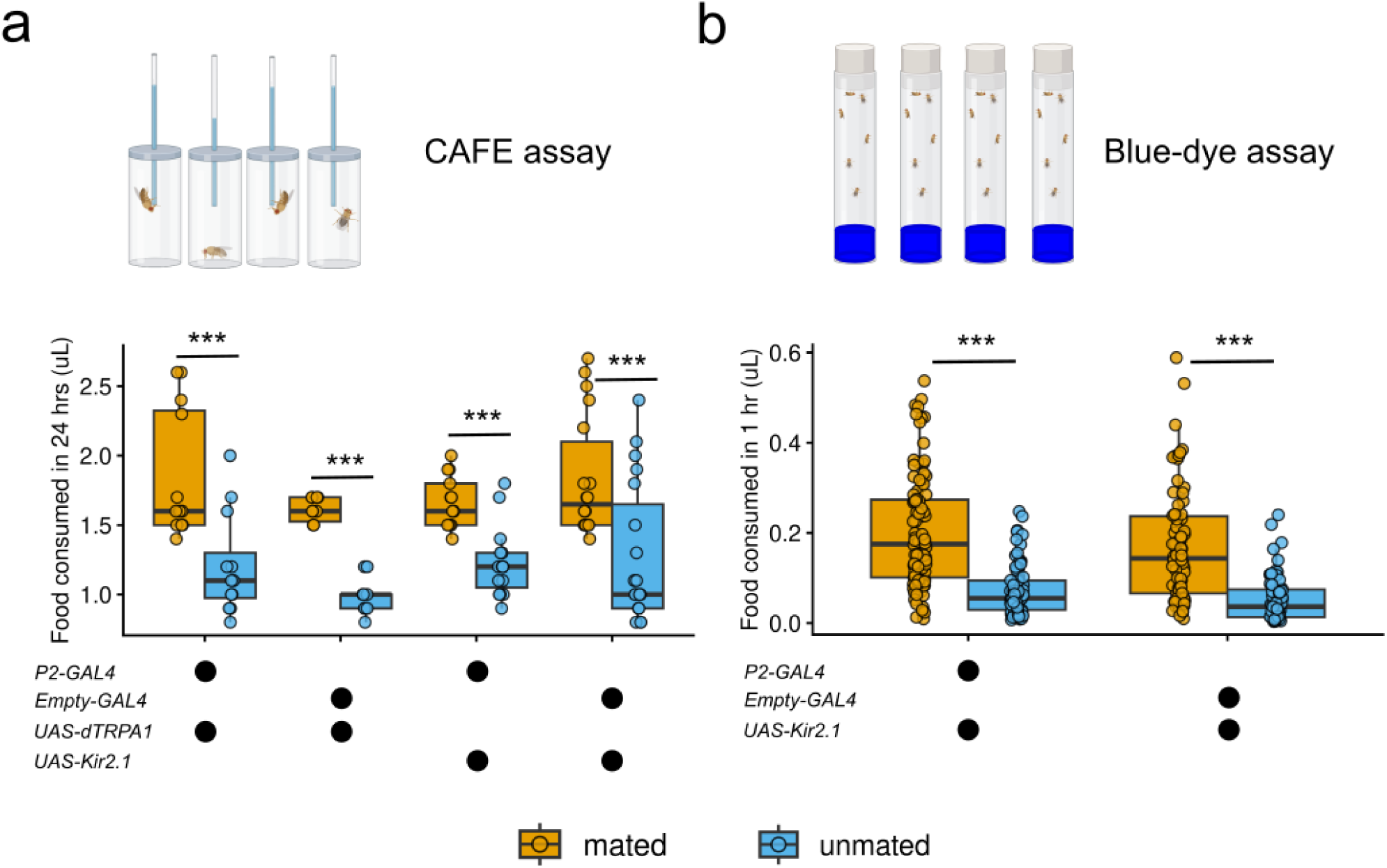
P2 neuron perturbations do not affect female post-mating food consumption. (a) Food consumed in 24 hours using CAFE assay in P2 neuron activated (*P2>dTRPA1*) mated (*n*=12), and unmated (*n*=12) females, their no GAL4 control (*EM>dTRPA1*) mated (*n*=10), and unmated (*n*=9) females, P2 silenced (*P2>Kir2.1*) mated (*n*=13), and unmated (*n*=15) females, and their no GAL4 control (*EM>Kir2.1*) mated (*n*=18) and unmated (*n*=19) females. (b) Food consumed in 1-hour using blue-dye extraction assay in P2 silenced (*P2>Kir2.1*) mated (*n*=98), and unmated (*n*=83) females, and their no GAL4 control (*EM>Kir2.1*) mated (*n*=74) and unmated (*n*=65) females. Amount of food consumed was compared statistically using two-way ANOVA followed by Sidak’s posthoc pairwise comparison. All boxplots depict the median, 25^th^ and 75^th^ percentiles, and whiskers represent 1.5 times the inter-quartile range. Dots in box and whisker plots represent sampled individuals. \*\*\**p* < 0.001. Figure not to scale, and created using biorender.com.

## Discussion

In *Drosophila melanogaster*, mating induces changes in female behaviors, with increased investment in feeding and egg laying, and reduced time sleeping during the day (Aigaki et al. 1991; Carvalho et al. 2006; Isaac et al. 2010; Garbe et al. 2016; Laturney et al. 2023). The male-transferred seminal fluid protein SP triggers these behavioral responses by acting on female mating-specific pC1 neurons in the brain (Yapici et al. 2008; Häsemeyer et al. 2009; Yang et al. 2009; Wang et al. 2020; Wang et al. 2021; Laturney et al. 2023). The neural circuitry underlying post-mating behaviors downstream of pC1 has only recently begun to be mapped. Here, we tested whether P2 neurons, which express sleep and feeding regulating NPF and innervate the sleep-regulating dorsal fan-shaped body in the central brain, mediate post-mating changes in sleep and feeding. Under the tested conditions, P2-neuron activation and silencing did not detectably alter the mating-associated reductions in daytime sleep or increases in food consumption.

We tested the role of P2 neurons on female post-mating sleep and feeding for two primary reasons. First, P2 neurons express neuropeptide F (NPF) and project to the dorsal fan-shaped body (dFB) (Lee et al. 2006; Shao et al. 2017; Li et al. 2021). The NPF-expressing dFB neurons regulate sleep homeostasis in flies (Jones et al. 2025) and modulate sleep in both socially isolated (Li et al. 2021) and post-mating contexts (Duhart et al. 2023). Distinct NPF-expressing dFB neurons regulate sleep-wake cycles by interacting with the fly’s internal hunger/satiety state (Chung et al. 2017), whereas activation of NPF neurons promotes feeding (Hergarden et al. 2012). A similar effect has been reported in bees, where experimental increase in NPF enhances foraging (Bestea et al. 2022). Evidence of NPF signaling in sleep and feeding regulation, together with its expression in P2 neurons, which project to sleep-regulating dFB, motivated us to test the role of P2 neurons in female post-mating changes in sleep and food consumption.

Second, P2 neurons are involved in social isolation-induced changes in sleep and food consumption (Li et al. 2021). The direction of behavioral change in the social isolation context is the same as that seen in females after mating (Carvalho et al. 2006; Isaac et al. 2010). Socially isolated flies with silenced P2 neurons showed sleep and feeding to levels comparable to those of group-housed flies, whereas acutely isolated flies with activated P2 neurons showed increased feeding and reduced daytime sleep (Li et al. 2021). However, in the mating contexts that we tested, these effects were not observed: mated females slept less during the day and consumed more food regardless of P2 neuron perturbation, similar to control mated females. These results suggest that distinct neurons may modulate similar behavioral changes in a context-dependent manner, and that P2 neurons may not be involved in the SP-influenced circuitry downstream of pC1 neurons.

Mating reduced female daytime sleep in all genotypes, but the effect lasted only 3-4 days in our study instead of ∼7 days as in Isaac et al. (2010). This shorter duration may be due to the sucrose-only food, which was intended to minimize interference from developing larvae interfering in the DAM assays. Consistent with previous reports (Yang et al. 2008; Yang et al. 2015; Garbe et al. 2016; Wang et al. 2020; Duhart et al. 2023), this food impeded egg-laying in all females in our studies, relative to standard food. While reduced egg laying and thus energy expenditure may account for the difference in duration of the post-mating sleep response we observed, differences in the genetic background between our lines and those used in the prior studies could also contribute.

Although mated females increase food intake after mating due to receipt of SP (Carvalho et al. 2006), there is the possibility that at least some of the underlying neural circuitry is nutrient-dependent. Mated females consume more sucrose (Laturney et al. 2023), salt (Walker et al. 2015), and protein (Walker et al. 2015; Liu et al. 2024) than unmated females. The increase in sucrose consumption by mated females is regulated by pCd-2 and Lgr3+ neuronal activity downstream of SP-sensing pC1 neurons (Laturney et al. 2023), whereas increase in protein consumption is mediated by octopamine (Walker et al. 2015) as well as ALK neurons expressing leucokinin (Liu et al. 2024). Because P2-neuron perturbation did not detectably alter the mating-associated increase in food consumption under our assay conditions, one or more of the neural circuits described above may contribute to post-mating feeding either downstream of, or independently from, SP-sensing pC1 neurons. It is also possible that neurons that innervate the gut could contribute to post-mating changes in feeding. The female midgut lengthens after mating, a process regulated by SP (Cognigni et al. 2011; White et al. 2021). The crop of mated females also enlarges after mating, a process regulated by myosuppressin neurons and their receptors, and involves ecdysone and bursicon α hormone signaling (Hadjieconomou et al. 2020). Although crop enlargement may accommodate increased food intake, its relationship to the SP-sensing neural circuitry is as yet unknown.

## Conclusion

Females of many animal species change their behavior after mating to increase fecundity and support offspring care. In the model organism *Drosophila melanogaster,* mated females increase food consumption and reduce daytime sleep. We tested the role of NPF-expressing P2 neurons on post-mating changes in sleep and feeding, because these neurons had been implicated in regulating these behaviors in a different social context. We found no evidence that P2 neurons contribute to the post-mating reduction in daytime sleep or the increase in food consumption. It is therefore possible that other neurons are responsible for modulating post-mating changes in female sleep and food consumption.

## Materials and Methods

### Drosophila stocks and crosses

All flies were maintained on a standard fly food medium (glucose (10% w/v), yeast (10% w/v), and agar (1.2% w/v)) at room temperature under a 12:12 light:dark cycle. *P2-GAL4* (*SS0020-P2-split-GAL4*), *UAS-Kir2.1* (*pJFRC49-10XUAS-IVS-eGFPKir2.1*), and *UAS-dTRPA1* (*UAS-dTRPA1/TM6b-brp/flp/Cyo*) flies were obtained from Wanhe Li (Li et al. 2021, Texas A&M University). *Empty-GAL4* (*Stable-split-empty-GAL4*) flies were obtained from Nilay Yapici (Cornell University). Crosses were used to generate *P2>dTRPA1* flies, whose P2 neurons were activated at 29°C, and *P2>Kir2.1* flies, whose P2 neurons were silenced. Control flies were the progeny of crosses between *Empty-GAL4* flies with *UAS-dTRPA1* flies and *UAS-Kir2.1* flies, respectively. We used *Canton-S* (CS) male flies for mass matings. Prior to the assays, P2 neuron expression in *P2-GAL4* flies was confirmed using *UAS-GFP.* Freshly dissected whole brains were imaged with an Echo Revolve RVL-100-G microscope (Supplementary Figure 1a, b, c). We also performed immunostaining with anti-GFP on fixed whole brains (Supplementary Figure 1d, e, f) as described in (Li et al. 2021). Virgin females were collected from these crosses and maintained in groups of 10-20 flies per vial. *CS* males were collected in parallel. All flies were 3-6 days old on the day of each experiment. One day prior to sleep and feeding assays, a subset of 3-6 day-old virgin females from the crosses described above were paired with *CS* males to obtain mated females.

### Sleep measurements

We measured locomotor activity in unmated and mated females using the Drosophila Activity Monitors (DAM) system (TriKinetics, Waltham, MA). We also conducted trials with females carrying the *UAS-Kir2.1* constructs at 25°C. Females were housed in a glass vial with sucrose-only food (5% 1:1 sucrose:agar). DAM assays were run for three consecutive days on the same females. Sleep profiles and durations were measured using Rethomics (Geissmann et al. 2019) in R. Sleep was defined as at least five minutes of uninterrupted inactivity. We compared total, daytime (ZT0-12), and nighttime (ZT12-24) sleep among mated and unmated females using ANOVA models that included genotype, mating status and their interaction, followed by pre-specified pairwise comparisons.

### Feeding measurements

We measured the food consumption per fly in unmated and mated females from the crosses described above using a capillary feeding (CAFE) assay modified from Murphy et al (2017). Briefly, we 3D-printed fly chambers, capillary gates, and the base as in (Murphy et al. 2017). We starved mated and unmated females overnight, allowing access to water. The following day, we set up a single fly per chamber, with 1% agar at the bottom of the chamber, and ∼5 µL of liquid food (5% sucrose, 5% yeast, ∼50 µL blue food dye) in 1-5 µL calibrated capillaries (Drummond Scientific). These capillaries were blocked with mineral oil on the other end to minimize evaporation of the liquid food. We set up 9 chambers without any flies as evaporation controls, and subtracted the food lost from the sample capillaries. We let the flies feed on the liquid food for 24 hours, and measured the food lost from the capillaries using vernier calipers. The amount of food consumed was analyzed with two-way ANOVA, treating genotype and mating status as fixed effects, followed by pairwise comparisons.

For *P2>Kir2.1* and its associated control, we also measured food consumption using a blue-dye assay. Mated and unmated females were starved overnight with access to water and then given food containing 2.5% (w/v) blue food dye (FD & C Blue Dye No. 1) as in Wong et al (2009). On the day of the experiment, flies were allowed to feed on blue-dyed food for 1 hour during the ZT0-ZT4 interval.

After feeding, each fly was individually washed in 1X PBS to remove excess blue dye adhering to the body, flash frozen in liquid nitrogen, decapitated and homogenized in 100µL 1X PBS using a variable-speed motor (Model S63C, Tri-R instruments) to extract the ingested blue-dyed food. The solution was centrifuged for 2 minutes, and 50 µL of the supernatant was used for spectrophotometric analysis.

Absorbance was measured at 629nm (Wong et al. 2009) on a Thermo scientific Varioskan Lux plate reader. The amount of food consumed by each fly was calculated using standard curves of serially diluted blue-dyed food, and compared with two-way ANOVA, treating genotype and mating status as fixed effects, followed by pairwise comparisons. We were unable to use this assay to measure feeding by *P2>dTRPA1* flies, since their genetically-matched control flies (*EM>dTRPA1*) did not show increased feeding post-mating (*t* = 1.42, df = 547, *p* = 0.15), in contrast to previous reports (Carvalho et al. 2006) and our own observations with the *UAS-Kir2.1* experimental and controls.

### Egg laying

We measured egg laying by mated flies of the four genotypes over three days on sucrose-only food (the food used for sleeping assays) and on standard fly food. Flies with *UAS-dTRPA1* constructs were housed at 29°C, whereas flies with *UAS-Kir2.1* constructs were housed at 25°C. Singly-mated flies were moved each day for three days to a new vial containing either sucrose-only or the standard food, and the eggs laid over each 24 hour period were counted. The numbers of eggs laid were compared across genotypes and food substrate using a two-way ANOVA with aligned rank transformation tests, with genotype and food substrate treated as fixed effects, followed by pairwise comparisons.

## Data availability

Raw data will be publicly made available upon manuscript publication

## Acknowledgments

We thank the members of the Clark and Wolfner labs and Nilay Yapici, Wanhe Li, Viveka Singh, and Sheetal Potdar for insights and discussions during this project and/or for comments on this manuscript. We thank Nilay Yapici and Wanhe Li for flies, Kate Scuderi and Vyom Mukerjee for assistance with data collection, Mehrnaz Afkhami with brain dissection, Emily L. Rivard for microscopy assistance, and John McCormick and Justin Galardi for spectrophotometry.

## Supplementary figures

**Supplementary Figure 1:**
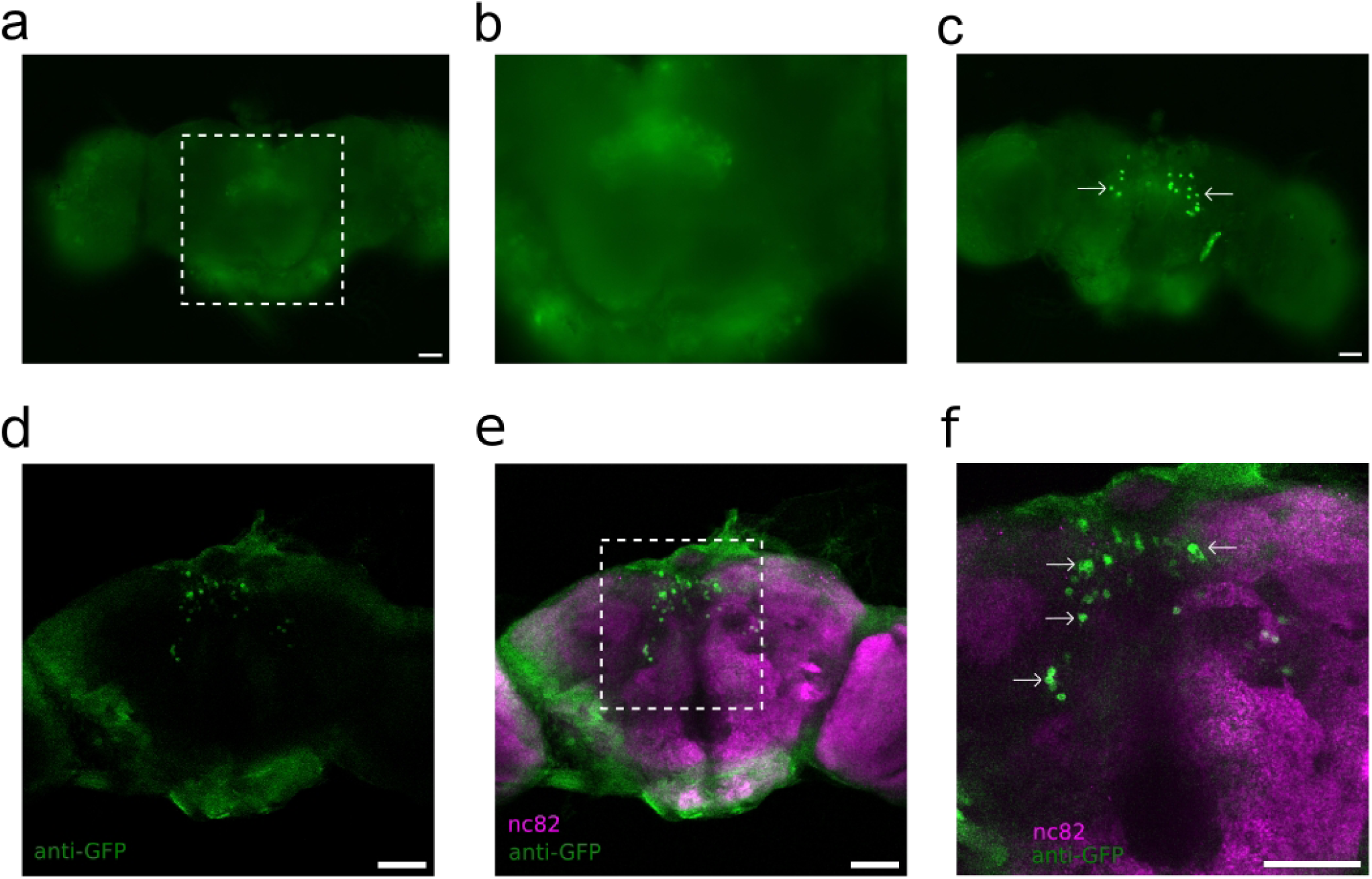
Confirming P2 neuronal expression of the P2-GAL4 driver line we used by UAS-GFP. a) The P2 neuron driven expression of GFP positive signal (*P2>GFP*) is localized in the fan layer of the central brain (white box), which is enlarged in (b). (c) The P2 neuron driven expression of GFP in the cell bodies, both as previously reported in (Shao et al. 2017; Li et al. 2021), and further confirmed by (d) anti-GFP antibody immunostain against (e) nc82 background (white box), enlarged in (f). Scale bars in (a,c) = 100µm, and in (d-f) = 50µm.

**Supplementary Figure 2:**
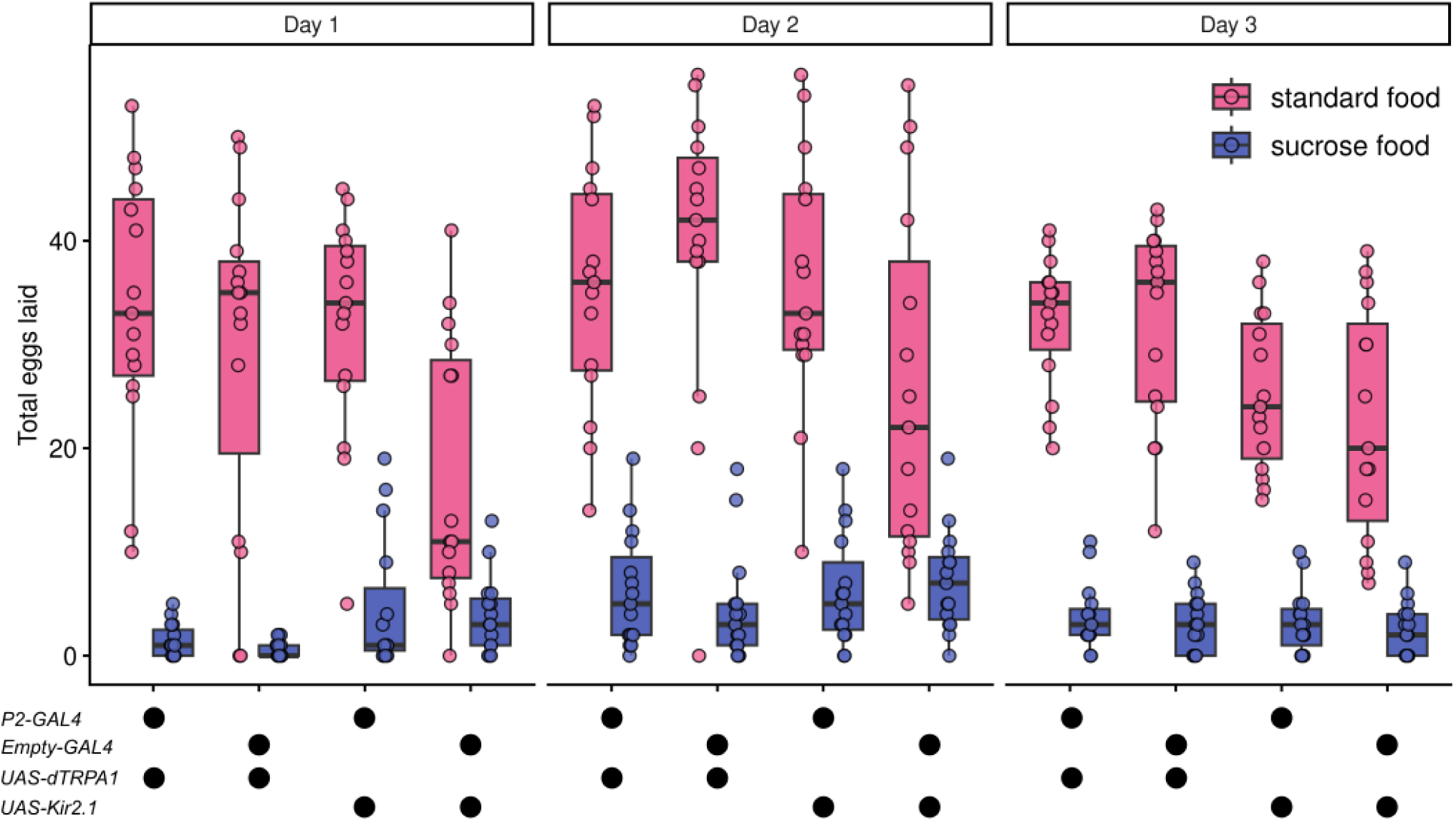
Irrespective of P2 neuron perturbation, mated females laid more eggs on standard food on all 3 days. The total number of eggs laid by mated females of different genotypes across three days on standard food that contains yeast, and on sucrose-only food that does not contain yeast. Mated females laid more eggs on standard food than sucrose-only food across all genotypes and days (*p* < 0.0001). Number of eggs laid were compared between the two food substrates within different genotypes and days using an aligned rank transformed ANOVA, followed by Tukey’s HSD pairwise comparison.

**Supplementary Figure 3:**
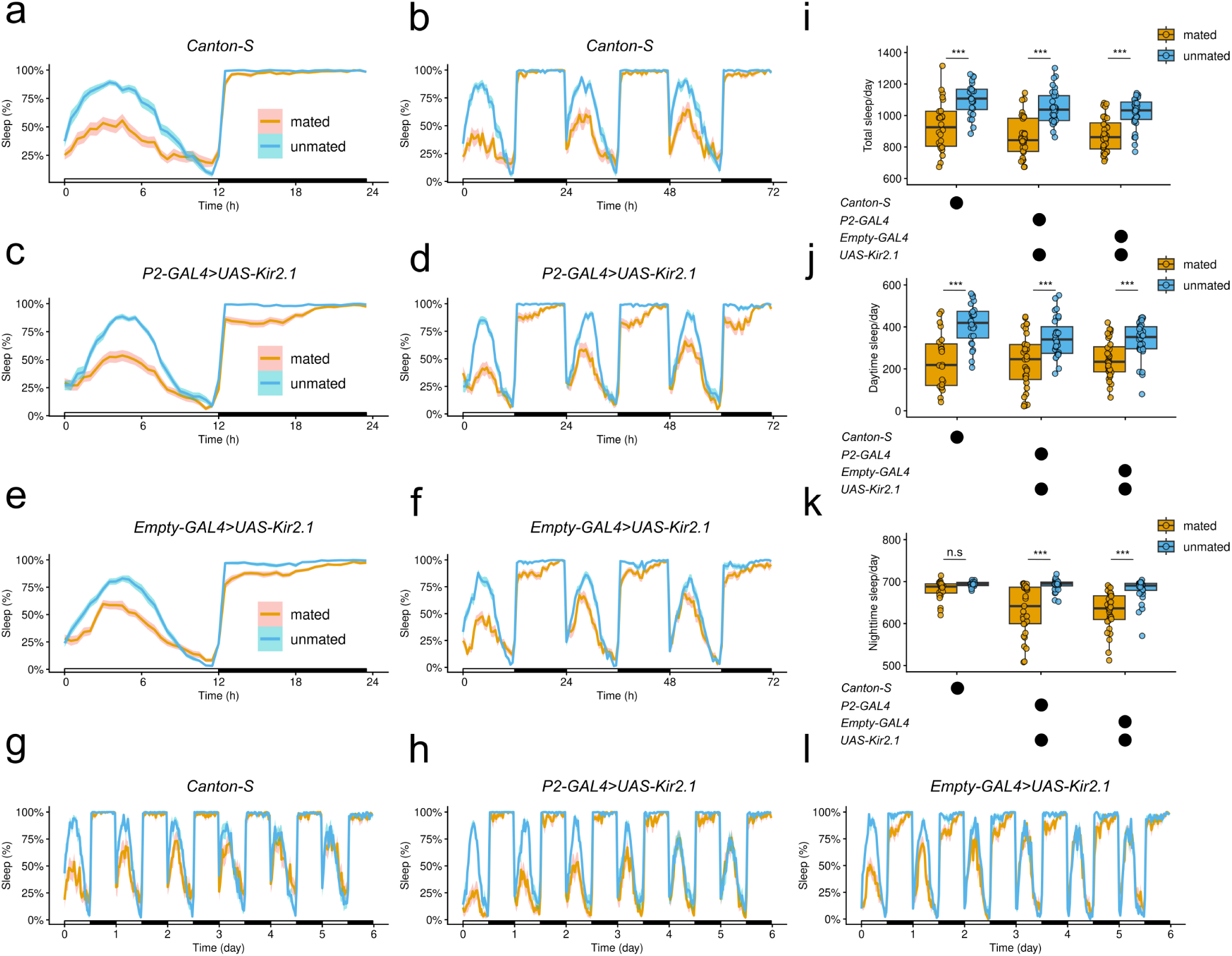
Altering P2 neuron activity does not modulate female post-mating daytime sleep at 25°C. Sleep profiles averaged across 3 L:D cycles in (a) *Canton-S* mated (n=24) and unmated (n=24) females, (c) P2 neuron silenced mated (n=32) and unmated (n=32) females, and its (e) no GAL4 control (*EM>Kir2.1*) mated (n=32) and unmated (n=32) females. (b), (d) and (f) depict daily sleep rhythms of the same genotypes and mating status as in (a),(c), and (e) respectively. Sleep for all flies were measured at 25°C in this figure. Mated females of all genotypes slept less compared to unmated females, averaged across L:D cycles, as measured by (i) total sleep; (j) daytime sleep; and (k) nighttime sleep, except for *Canton-S*. In one of the trials, we measured sleep across 6 L:D cycles, and mated females of (g) *Canton-S*, (h) *P2 silenced*, and (l) no GAL4 control revert back to unmated like sleep states by day 4 (n=14-15 flies/genotype/mating status). Sleep was statistically compared using two-way ANOVA followed by Sidak’s posthoc pairwise comparison. All boxplots depict median, 25^th^ and 75^th^ percentiles, and whiskers represent 1.5X inter-quartile range. Dots in box and whisker plots represent sampled individuals. n.s. = not significant, \**p* < 0.05, \*\**p* < 0.01, \*\*\**p* < 0.001.

**Supplementary Figure 4:**
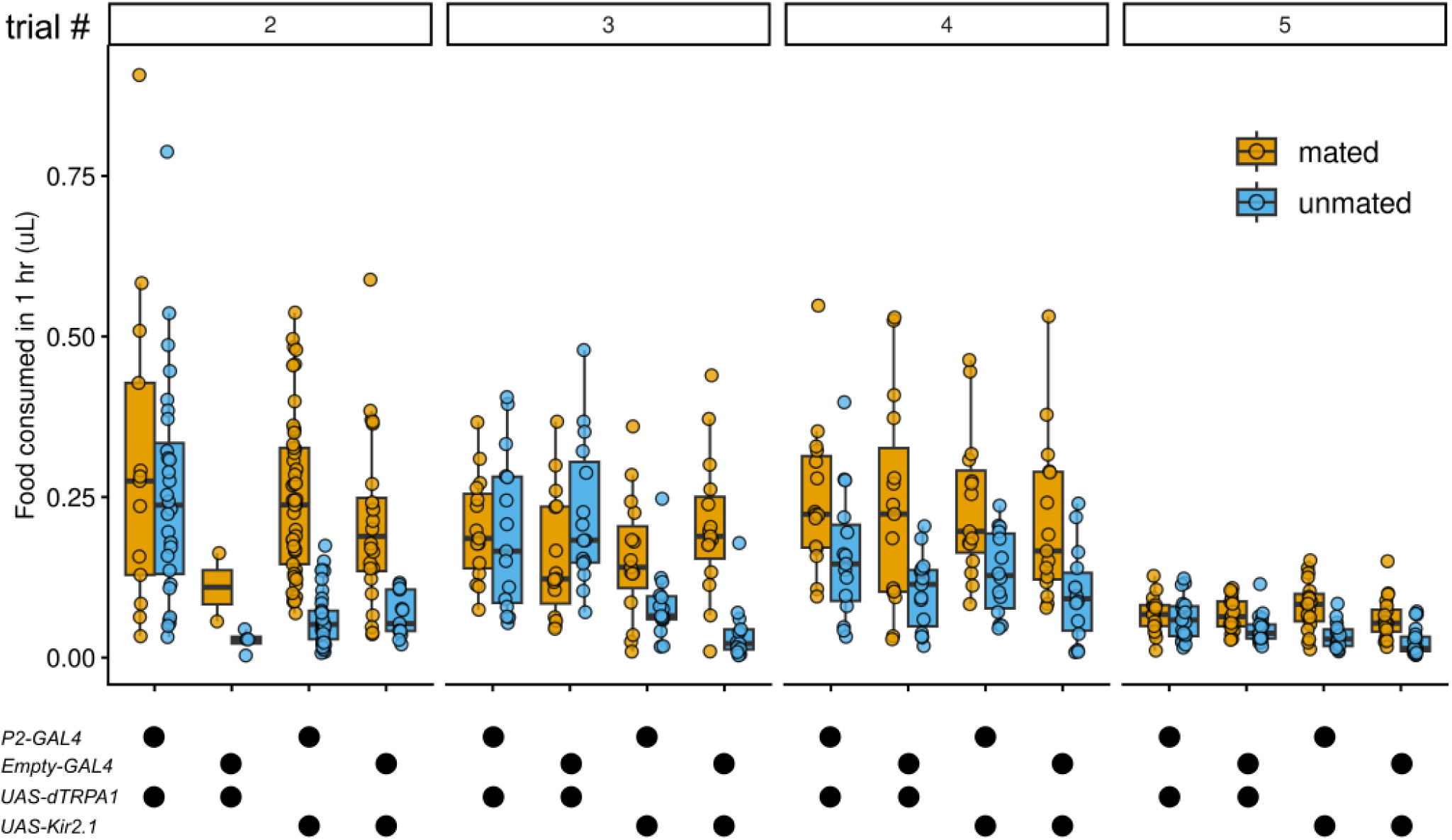
Food consumed by mated and unmated females varied across different trials. The amount of food consumed by mated and unmated females of the four genotypes were measured using blue dye extraction assay across four trials, and we found a significant effect of trial on the amount of food consumed (ANOVA, p<0.0001). In all trials, flies with the *UAS-Kir2.1* constructs ate more compared to their unmated counterparts.

